# Thermal acclimation mitigates cold-induced paracellular leak from the *Drosophila* gut

**DOI:** 10.1101/136895

**Authors:** Heath A. MacMillan, Gil Yerushalmi, Sima Jonusaite, Scott P. Kelly, Andrew Donini

## Abstract

When chilled to temperatures below their critical thermal minimum, chill susceptible insects can suffer tissue damage and die. The mechanisms that cause this chilling injury are not well understood but a growing body of evidence suggests that a loss of ion and water homeostasis in the cold leads to hemolymph hyperkalemia that depolarizes cells, leading to cell death. The apparent root of this cascade of issues is the net leak of osmolytes down their concentration gradients in the cold. Many insects, however, are capable of adjusting their thermal physiology, and cold-acclimated *Drosophila* can maintain homeostasis and avoid chilling injury better than warm-acclimated flies. Here, we test whether the cold-induced loss of osmotic homeostasis is associated with a loss of epithelial barrier function in *Drosophila*, and provide the first evidence of cold-induced epithelial barrier failure in an invertebrate. Flies exposed to 0° C had increased rates of paracellular leak through the gut epithelia, but cold acclimation reduced paracellular permeability, both before and during cold stress, and improved cold tolerance. This adjustment in barrier function was associated with changes in the abundance of select septate junction proteins and the appearance of a tortuous ultrastructure in subapical intercellular regions of contact between adjacent midgut epithelial cells. Thus, cold causes paracellular leak in a chill susceptible insect and cold acclimation can mitigate this effect, at least partly through changes in the composition and structure of transepithelial barriers.

**Summary Statement:** Chilling disrupts barrier function of the gut of flies and cold acclimation can mitigate this problem through changes in paracellular occluding junctions.

## Introduction

The majority of animal species are insects, and the majority of insects are chill susceptible. By definition, chill susceptible insects suffer from non-freezing chilling injury at temperatures below species-specific physiological thresholds^1,2^. When chilled, chill susceptible insects cross their critical thermal minimum (CT_min_) and enter a state of complete paralysis termed chill coma^3^. If they are subsequently moved to warmer temperatures, insects will recover from chill coma with few apparent injuries, and the time required to recover to a standing position is termed chill coma recovery time (CCRT)^4^. Prolonged exposure to low temperatures, however, causes chill susceptible insects to suffer tissue damage and die^1^. The CT_min_, CCRT and survival at low temperatures are all useful metrics of chilling tolerance, and all appear to be mediated by related but distinct physiological mechanisms^1^.

One rapid effect of chilling is a suppression of ion-motive ATPase activity. In excitable cells, an immediate reduction in active ion transport with cooling reduces the electrogenic contribution of ion transport to cell membrane potential^5–7^. In the muscles, this loss of cell potential may strongly contribute to hyper-excitability and ensuing electrical silence at the onset of chill coma^5,6,8^. In the nervous system, a rapid and reversible surge of extracellular K^+^ (an immediate consequence of reduced Na^+^/K^+^-ATPase activity in glial cells) further depolarizes nerve cells near the chill coma onset temperature^9,10^. During prolonged exposure to low temperatures, chronic suppression of active transport relative to passive leak rates can cause a progressive loss of ionic and osmotic balance in a variety of model insects, including representatives of Diptera, Orthoptera, Hemiptera, and Lepidoptera^1,11^. These chronic effects of chilling are most evident at the insect gut and Malpighian tubule epithelia, where high rates of ion transport normally serve to regulate organismal ion and water homeostasis^12^. In the cold, net leak of Na^+^ (a common major osmolyte) down its concentration gradient drives a parallel migration of water into the gut lumen, dehydrating the hemocoel^13,14^. This loss of hemolymph water, in some cases combined with a net leak of K^+^ from the tissues into the hemolymph^14^, causes a rise in hemolymph concentration of K^+^, which depolarizes cells, leading to muscle cell death^15,16^.

Chilling can thus have profound effects on ion and water balance, and the ultimate consequences of a loss of homeostasis can be injury and death. Chill tolerance can, however, vary widely as a result of the evolution of tolerance evident under common-garden conditions^17,18^ or the evolution of long-or short-term phenotypic plasticity in chill tolerance^19–21^. *Drosophila* represent an ideal model system for studying variation in chill tolerance among and within species. Species of this genus vary in CT_min_ by more than 11° C^17^, a range that can also be mirrored within a single cosmopolitan species (*D. melanogaster*) by combining developmental and adult thermal acclimation with rapid heat-and cold-hardening^22^. Cold acclimation reduces the effects of chilling on ion and water homeostasis, such that tissue integrity is better preserved during chronic cold stress. Cold-acclimated *D. melanogaster*, for example, constitutively maintain lower Na^+^ concentration gradients, and better maintain hemolymph water and [K^+^] balance during a cold stress^23^. Taken together, this body of evidence suggests that chilling injury can result from an inability to maintain rates of ion transport relative to leak, and that improvements in chill tolerance can be driven by thermal plasticity of ionomotive epithelia.

Naturally, the impacts of chilling on transcellular ion and water flux in the Malpighian tubules and gut epithelia are under active investigation^14,24^. Solutes, however, can cross epithelial barriers by either transcellular or paracellular routes^25^. In insects, paracellular barriers are established by the septate junctions (SJs); paracellular occluding junctions that are functionally analogous to the vertebrate tight junction (TJ) complex^26,27^. SJs are multi-protein complexes and in cross-sectional view appear as ladder-like structures between adjacent cells with septa spanning a 15-20 nm intercellular space^27^. In insects, two morphological variants of SJs exist based on tangential sections and these are the pleated SJs and the smooth SJs. The septa of the pleated SJs appear as regular undulating rows and this type of SJ is found principally in the ectodermally-derived tissues (e.g. trachea, foregut and hindgut)^27^. The septa in the smooth SJs show regularly spaced linear lines and these junctions are located in the endodermally-derived tissues (e.g. midgut and Malpighian tubules)^27^.

SJs can form selective and dynamic molecular barriers to solute diffusion. For example, in the midgut epithelium of a caterpillar (*Bombyx mori*), low-resistance junctions are lined with negative charge, facilitating selective reabsorption of cations^28^. By contrast, anions (Cl^-^) follow cations passively through the paracellular pathway of the Malpighian tubules in a mosquito (*Aedes aegypti*), where rapid regulation of paracellular permeability is achieved by leucokinin signalling^29,30^. Under favorable conditions, SJs in the gut epithelia effectively separate the organism from the outside environment (i.e. gut lumen), but it has been noted that barrier function is compromised in *Drosophila* under conditions that lead to death, such as advanced age^31^, or following traumatic brain injury^32^. Such a loss of gut barrier function allows for large polar molecules like glucose and even the intestinal microbiota to leak into the hemolymph^32^.

Because exposure to low temperature has been documented to suppress active solute transport across insect epithelia resulting in a progressive loss of ionoregulatory homeostasis, an open question is how does chilling impact the molecular physiology of SJs and the functional dynamics of the paracellular pathway? Indeed, given that cold acclimation mitigates the deleterious effects of chilling on insect salt and water balance, and yet is associated with decreased (rather than increased, as might be predicted) activity of ion-motive ATPases in *D. melanogaster*^33^, it can be hypothesized that SJ plasticity contributes significantly to cold tolerance plasticity. This idea is supported by recent observation that cold acclimation leads to upregulation of approximately 80% of genes encoding currently known or putative *Drosophila* SJ proteins^34^ (see Table S1).

## Materials and Methods

### Animal husbandry

The line of *Drosophila melanogaster* (Diptera: Drosophilidae) used in this study were derived from 35 isofemale lines collected in London and Niagara On The Lake, Ontario, Canada in 2007 as previously described^35^. Flies were reared from egg to adult in 200 mL bottles containing 50 mL of Bloomington *Drosophila* medium^36^ at constant 25° C and a 14 h:10 h light:dark cycle. Adult flies were transferred to fresh bottles for 2 h to oviposit before being removed in order to ensure a rearing density of ∼100 eggs per bottle. Adults (both males and females) were collected on their day of emergence and transferred to 40 mL vials containing 7 mL of fresh diet at a density of ∼40 flies per vial. Half of the adult flies were maintained at 25° C (control, warm-acclimated conditions) and the other half were transferred to a separate incubator at 10° C with a 10 h:14 h light:dark cycle for cold acclimation conditions. Female flies were sampled for experiments seven days after final ecdysis and males were discarded. Thus, all experiments were completed on seven day old mated females.

### Chilling tolerance

We quantified the cold tolerance of control and cold-acclimated animals by determining the critical thermal minimum (CT_min_), chill coma recovery time (CCRT), and survival at 0° C of experimental animals. The CT_min_ was measured as previously described^8^. Briefly, flies were transferred to 4 mL glass vials and placed in an aluminum rack that was suspended in ethylene glycol. The glycol was circulated by a refrigerated circulating bath (MX7LL, VWR International, Mississauga, Canada) that was preset to 20° C. The temperature was then lowered at a rate of 0.15° C min^-1^ and the flies were continuously observed. Temperature was confirmed and recorded using a TC-08 interface equipped with two type-K thermocouples (Pico Technology, St. Neots, UK). The CT_min_ was quantified as the temperature at which no movement was observed from a fly when the vial was tapped with a probe.

We measured CCRT as previously described^23^. Flies were transferred individually into 4 mL glass vials as above but the vials were then placed in a mixture of ice and water and thereby maintained at 0° C for 6h. The vials were then quickly removed to the laboratory bench at 23° C and the flies were observed. Recovery time was scored as the time required for a fly to spontaneously right itself and stand on all six legs without any form of stimulation.

Pilot experiments were used to select appropriate exposure times at 0° C to quantify low temperature survival in warm-and cold-acclimated flies following previously described methods^8^. Warm-acclimated flies were exposed to 0° C for up to 36 h and cold-acclimated flies were exposed to 0° C for up to 110 h. For each time point, groups of 7-12 flies were placed into 3-6 empty 40 mL vials, and the vials were suspended in an ice-water mixture as above. After removal from the cold the flies (still in chill coma) were transferred to new vials containing 3 mL of food. The vials with the recovering flies were left on their side at the respective acclimation temperature for 24 h before survival was assessed. Flies then able to stand and walk were counted as alive, and flies that were either not moving or moving but unable to stand and walk were counted as dead.

### FITC-dextran gut leak assay

Fluorescently-labelled dextran is commonly used to quantify paracellular permeability in vertebrate models (e.g. ^37,38^). To test for dextran leak from the *Drosophila* gut we mixed 50 μL of a 2.5% w/v solution of FITC-dextran (3-5 kDa, Sigma Aldrich) in water with 10 mg of dry active brewer’s yeast. The fluorescently labelled yeast paste was placed into a small feeding tray inside an otherwise empty 40 mL vial along with female flies (∼20 per vial). The vial was placed along with a kimwipe saturated with water (to maintain high humidity) inside a clear plastic box which was then put back into the 10° C or 25° C incubator for 24h. Early trials confirmed that all of the warm acclimated flies (12/12) and the majority of the cold-acclimated flies (11/12) consumed the FITC-dextran yeast mix within 4h, so 24h was chosen to ensure that both groups had the mixture in their gut for approximately the same amount of time before sampling. Diet can impact *Drosophila* cold tolerance (e.g. ^39^), so it was confirmed that feeding on the yeast and dextran mixture did not strongly influence CT_min_ nor CCRT (see Fig. S1). Subsequently the concentration of FITC-dextran in the hemolymph of control flies (fed FITC-dextran but not cold exposed) and those that received 6h or 24h at 0° C in an ice-water mixture was measured. Hemolymph was collected into small rectangular glass capillaries with a 20 μm path length (VitroTubes; VitroCom, Mountain Lakes, NJ, USA) and analyzed by confocal microscopy as described for primary urine droplets by Leader and O’Donnell^40^. The capillary was used to collect the hemolymph droplet emerging from an ablated fly antenna under air pressure^41^ and the ends of the capillary were carefully dipped into paraffin oil to prevent evaporation. The capillary containing the sample was then placed on a glass slide and viewed using a Fluoview 300 confocal laser scanning microscope (Olympus America, Center Valley, CA, USA) with a 488 nm excitation laser and 535-565 bandpass emission filter. Laser power was minimized (10% of maximal) to avoid photobleaching. Two different objective lenses (20× and 40×) and sets of microscope settings were selected based on early trials and used depending on the apparent concentration of FITC-dextran in the sample. A z-stack of images was taken for each sample, covering ∼30 μm around the center of the capillary. Image analysis was completed in FIJI (a distribution of imageJ)^42^. For each set of images, four areas (each approximately 200 × 200 pixels) within the capillary and one outside of the capillary were selected. The fluorescence intensity of each region was averaged and subtracted from the background (outside the capillary) and concentration was determined using the maximal fluorescence value (taken at the optical centre of the tube) by reference to standard curves (shown in Fig. S2).

### Tissue damage

To examine whether leak across the gut epithelia in the cold was likely to be driven by the accumulation of chilling injury (i.e. cell death), we used a Live/Dead sperm viability kit (Thermo Fischer Scientific, Waltham, MA, USA) following previously described methods^43^ with some modifications. Warm-and cold-acclimated flies were exposed to 6, 24 or 48 h at 0° C as described for the survival assay. Immediately upon removal from the cold each fly was dissected using micro forceps in *Drosophila* saline to remove the entire gut without directly touching the gut tissue. Each gut was individually placed on a slide in 10 μL of Sybr-14 (25 μM) in *Drosophila* saline for 10 min, after which 10 μL of 72 μM propridium iodide was added for a further 10 min. A coverslip was gently placed over the tissue and the gut was imaged with a 20× objective lens using a fluorescence microscope (Nikon Eclipse Ti-S, Nikon Instruments Inc., Minato, Japan) in five regions (the crop, anterior midgut, posterior midgut, ileum, and rectum). For Sybr green, we used a filter set intended for FITC (ex: 488, em: 519) which is very close to the Sybr 14 excitation and emission maxima (ex: 480, em: 516). For propridium iodide (ex: 525, em: 617), we used a filter set intended for Texas red (ex: 560, em: 610). For each region of interest a z-stack of images was converted into a single focused image by the microscope software (NIS Elements, Nikon). Image analysis was completed in FIJI as described previously ^15^. Briefly, the green (Sybr-14 stained, live cells) and red (propridium iodide stained, red cells) channels were separated and converted to greyscale before a threshold was applied to each image to highlight the green or red area of the image. Proportions of red and green particles within each image (i.e. cell counts) were quantified, as were the proportions of red and green area within the image. These two approaches yield slightly different estimates of damage because propridium iodide binds DNA, and DNA is degraded during cell death^15,16^. Thus, we averaged the two methods of analysis to obtain a proportion of injured tissue for each image.

### Immunoblotting

To quantify the relative abundance of known proteins of *Drosophila* SJs, whole warm-and cold-acclimated flies were counted into 1.7 mL microcentrifuge tubes (10 flies per sample) and snap frozen in liquid nitrogen before being stored at -80° C. Flies were later homogenized in a buffer (50 mM Tris–HCl, 150 mM NaCl, 1 % sodium deoxycholate, 1 % Triton-X-100, 0.1 % SDS, 1 mM PMSF and 1:200 protease inhibitor cocktail [Sigma Aldrich], pH 7.5) by grinding with a plastic pestle prior to sonication on ice (4 × 10 s with 30 s rests). Homogenized samples were centrifuged at 12,000 × *g* for 10 min and aliquoted before protein content was determined in a subsample using a Bradford assay (Sigma-Aldrich). For electrophoretic separation by SDS-PAGE, samples containing 10 μg of total protein were added to a 6 × loading buffer (360 mM Tris–HCl, 12 % (w/v) SDS, 30 % glycerol, 600 mM dithiothreitol and 0.03 % (w/v) bromophenol blue) and separated following previously described methods^44^. Primary antibodies and antibody dilution factors used are listed in Table S2. Clarity^™^ Western ECL substrate (Bio-Rad, Hercules, CA, USA) was used to visualize antigen activity with a Gel Doc XR+ system (Bio-Rad). After probing and imaging, membranes were stained for total protein with coomassie blue as a loading control as described by Welinder and Ekblad^45^. Antigen expression was normalized relative to the loading control, and for Kune, two immunoreactive bands were analyzed independently.

### Electron microscopy

Warm-and cold-acclimated female flies were dissected in *Drosophila* saline to carefully remove the entire gut. Gut samples for electron microscopy were fixed in 2% (v/v) glutaraldehyde in 0.1M sodium cacodylate, rinsed once in the same buffer, and post-fixed in 1% osmium tetroxide. The samples were dehydrated in a graded ethanol series followed by propylene oxide. The entire gut was then viewed using a dissecting microscope and cut at the midgut-hindgut junction before samples were embedded in Quetol-Spurr resin. Cross-sections 90 nm thick were cut into the posterior end of the posterior midgut on an Leica Ultracut ultramicrotome before being mounted on grids and stained with uranyl acetate and lead citrate. Posterior midgut sections were imaged at 25,000 × magnification in a Tecnai 20 transmission electron microscope (FEI, Hillsboro, OR, USA). Images of cell-cell contact in the subapical region of posterior midgut cells (n=4-7 cell-cell regions of 3 biological replicates per acclimation group) were captured and analyzed in Fiji. Briefly, a 2000 × 2000 pixel subsample of each image was isolated and the line tool was used to draw over any regions of cell-cell contact visible within the area. From this tracing the total length of visible cell-cell contact was determined. The same approach was used to determine the length of visible SJs, which we identified as dark regions of contact with a characteristic “ladder-like” appearance. This entire analysis was completed by an experimenter who was blind to the identity of the images.

### Data analysis

All data analysis was completed in R version 3.3.2^46^. We tested for effects of acclimation temperature on CT_min_ and CCRT using Welch two-sample t-tests, and survival data were analyzed using a generalized linear model (glm) with a binomial error distribution and a logit link function. The effects of acclimation and cold exposure on the accumulation of FITC-dextran in the hemolymph were analyzed using an ANCOVA on log_10_ concentrations. We tested for effects of acclimation temperature and cold exposure duration on levels of tissue damage independently in each section of the gut using ANCOVAs, followed by pairwise t-tests which were corrected for false discovery using the method of Benjamini and Hochberg^47^. Abundance of SJ proteins in warm and cold acclimated flies were compared using Welch two-sample t-tests on log-transformed relative density. The length of visible cell-cell contact and SJs within images were compared using a nested ANOVA, with individual junctions nested within biological replicates.

## Results

### Chilling tolerance

Cold acclimation led to improvements in chilling tolerance based on three endpoints; CT_min_, CCRT and survival. First, adult female flies acclimated to 10° C had a CT_min_ of 5.3 ± 0.1° C, 3° C lower than those acclimated to 25° C (8.3 ± 0.1° C; t_35_=22.0, *P* < 0.001; Fig. 1A). Second, cold-acclimation significantly reduced CCRT after 6h at 0° C (t_27_=7.2, *P* < 0.001); cold-acclimated flies recovered in 14.6 ± 0.8 min, nearly 50% faster than warm acclimated flies that recovered in 27.5 ± 1.6 min (Fig. 1B). Finally, flies acclimated to 10° C survived exposure to 0° C for significantly longer than those acclimated to 25° C (z_910_=9.35, *P* < 0.001; Fig. 1C); the time required to cause 50% mortality at 0° C (Lt_50_) was roughly doubled in cold-acclimated flies (59.5 ± 1.9 h) relative to warm-acclimated flies (29.2 ± 0.6 h).

**Figure 1.**
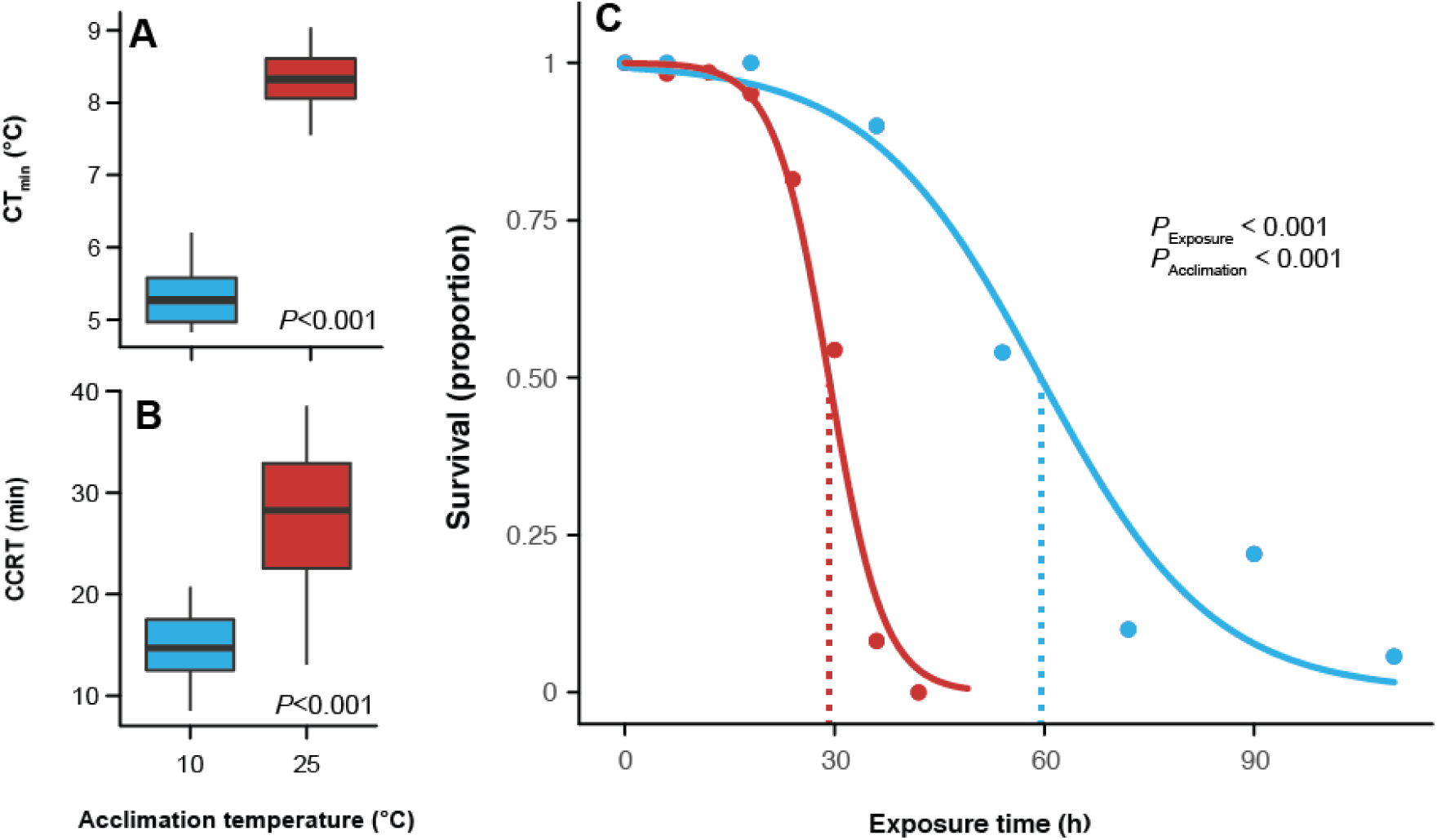
Cold acclimation improves chilling tolerance of female *Drosophila melanogaster*. The critical thermal minimum (CT_min_, A), chill coma recovery time following 6h at 0° C (CCRT, B) and survival following exposure at 0° C (expressed as a proportion of flies surviving, C) of warm-(red) and cold-acclimated (blue) female *D. melanogaster*. *P*-values in panels A and B denote significant differences in the CT_min_ and CCRT, respectively. *P*-values in panel C denote main effects of a binomial logistic regression, which was used to determine L_t_50 values (dotted lines).

### FITC-Dextran leak assay

To determine whether chilling specifically induces paracellular leak across the gut epithelia in *Drosophila*, we quantified the appearance of FITC-dextran (which can only pass via the paracellular pathway) in the hemolymph of flies fed a mixture of FITC-dextran in yeast. Flies fed the FITC-dextran yeast had the mixture distributed throughout their entire gut lumen (Fig. 2A). Examples of hemolymph samples collected from control 25° C-and 10° C-acclimated flies are shown in Fig. 2B. We found a significant interactive effect of acclimation temperature and cold exposure on the appearance of FITC-dextran in the hemolymph (F_1,41_=5.12, *P* < 0.03), as well as main effects of acclimation temperature (F_1,41_=140.0, *P* < 0.001) and cold exposure duration (F_1,41_=114.9, *P* < 0.001). Warm-acclimated flies had approximately 7.6-fold higher concentrations of FITC-dextran in their hemolymph (27.2 ± 4.8 ng μl^-1^) than cold-acclimated flies (3.6 ± 0.6 ng μl^-1^; Fig. 2C) before any cold exposure was experienced. Cold exposure caused the concentration of FITC-dextran in the hemolymph of warm-acclimated flies to increase by 26-fold after 24h at 0° C (727.4 ± 128.9 ng μl^-1^), while cold-acclimated flies reached concentrations about 10.5-fold higher (38.7 ± 9.2 ng μl^-1^) than before the cold exposure (Fig. 2C). Warm-acclimated flies that were exposed to 0° C for 16h, allowed to recover for 6h at 25° C, and confirmed to be uninjured also had high levels of FITC-dextran in their hemolymph (640.1 ±195.9 ng μl^-1^; Fig. 2C).

**Figure 2.**
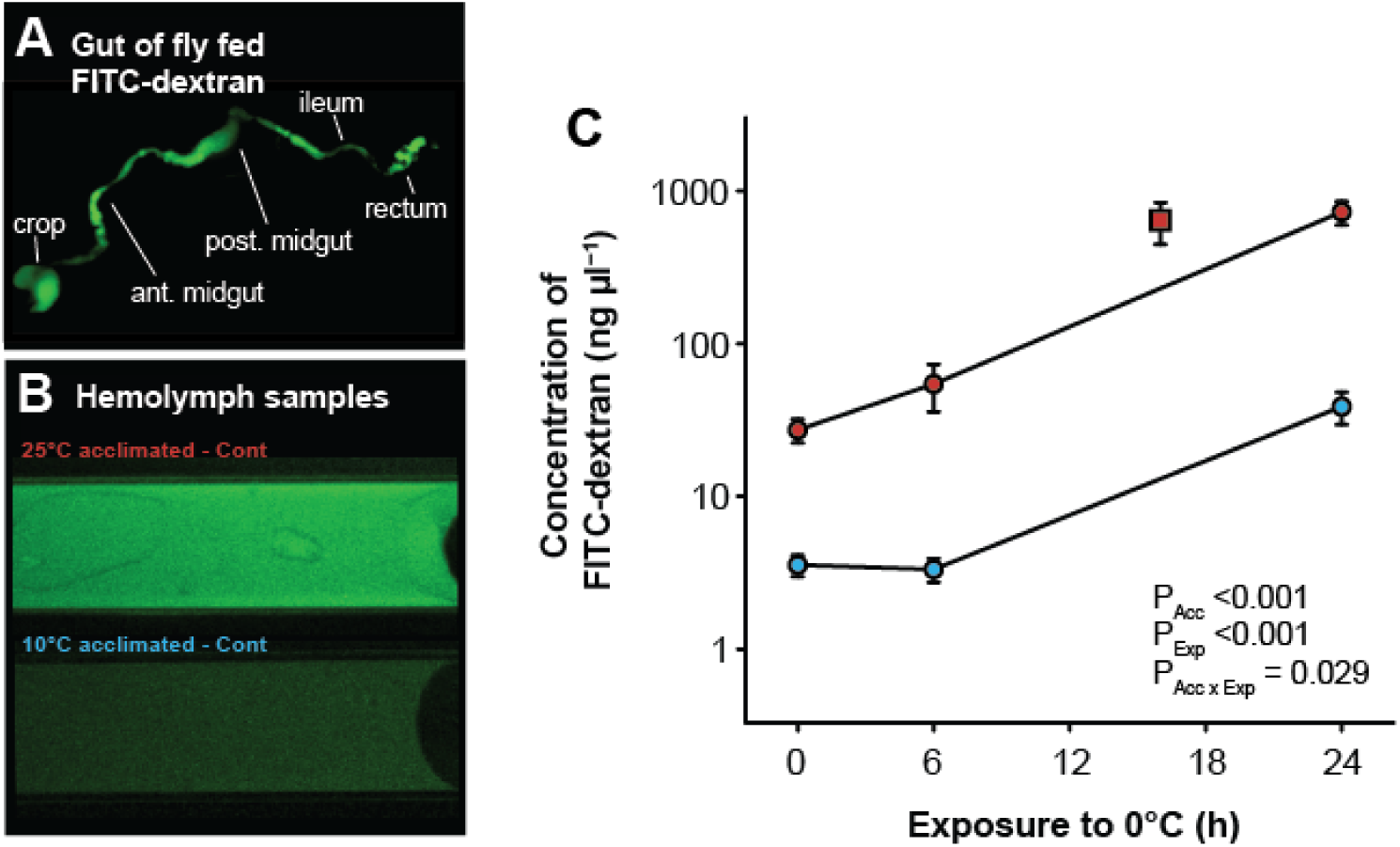
Cold acclimation preserves paracellular barrier function before and during chilling stress. A) The gut of a cold-acclimated female *Drosophila melanogaster* adult after feeding on FITC-dextran for 24h. Note that FITC-dextran is distributed through the entire gut. B) representative samples of hemolymph taken from warm-and cold-acclimated flies prior to any cold exposure. Images were taken with identical microscope settings. C) Mean ± sem concentrations of FITC-dextran (log_10_ scale) in the hemolymph of warm-(red) and cold-acclimated (blue) flies (n= 5-9 samples per group × exposure duration) before and following exposure to 0° C for up to 24h. Warm-acclimated flies had higher levels of FITC-dextran leak into the hemolymph prior to cold stress, and chilling caused FITC-dextran to leak at greater rates into the hemolymph of both acclimation groups. The single red square denotes FITC-dextran concentrations in the hemolymph of warm-acclimated flies which were exposed to 0° C for 16h, confirmed to have suffered no chilling injury after 6h of recovery at 25° C, and then sampled. This group had high concentrations of FITC-dextran despite not being injured, suggesting that leak is associated with compromised paracellular pathway integrity and not tissue damage in moribund animals.

### Intestinal damage assay

To determine whether chilling causes injury to the gut epithelia, and confirm that leak from the gut lumen was not a consequence of cell death occurring before leak was measured, we quantified levels of tissue damage in the gut epithelia of warm-and cold-acclimated flies before and following exposure to 0° C (Fig. 3A-T). Overall, damage to the gut occurred primarily in the midgut of warm-acclimated flies, and only at cold exposure durations beyond those observed to cause leak. In the crop, there was a significant effect of cold exposure duration on the proportion of cells labeled as “dead” (red) by propridium iodide (F_1,44_= 29.7, *P* < 0.001), but no significant effect of acclimation temperature (F_1,44_= 0.1, *P* = 0.975). Both warm and cold acclimated flies suffered some damage to the crop during chilling, but only after 48h at 0° C (Fig. 3A). There were significant interactions in the effects of acclimation temperature and cold exposure duration on tissue damage in both the anterior (F_1,46_=50.6, *P* < 0.001) and posterior midgut (F_1,46_=64.8, *P* < 0.001); chilling damaged both regions of the midgut, but only in warm-acclimated flies, and only after 48h at 0° C (Fig. 3B,C). No increase in damage was observed in the ileum of either warm-or cold-acclimated flies with increasing duration of cold exposure (F_1,46_ = 3.4, *P* = 0.071), nor was there any main effect of acclimation temperature (F_1,46_ = 2.7, *P* = 0.106; Fig. 3D). There was a marginally significant effect of cold exposure duration on tissue damage in the rectum (F_1,46_ = 4.2, *P* = 0.045), and the acclimation groups did not differ in the level of damage to the rectum (F_1,46_ = 0.4, *P* = 0.515). This tendency for the rectum to accumulate damage was small, and pairwise t-tests revealed no specific cold exposure duration that caused more damage than that observed in control animals (Fig. 3E).

**Figure 3.**
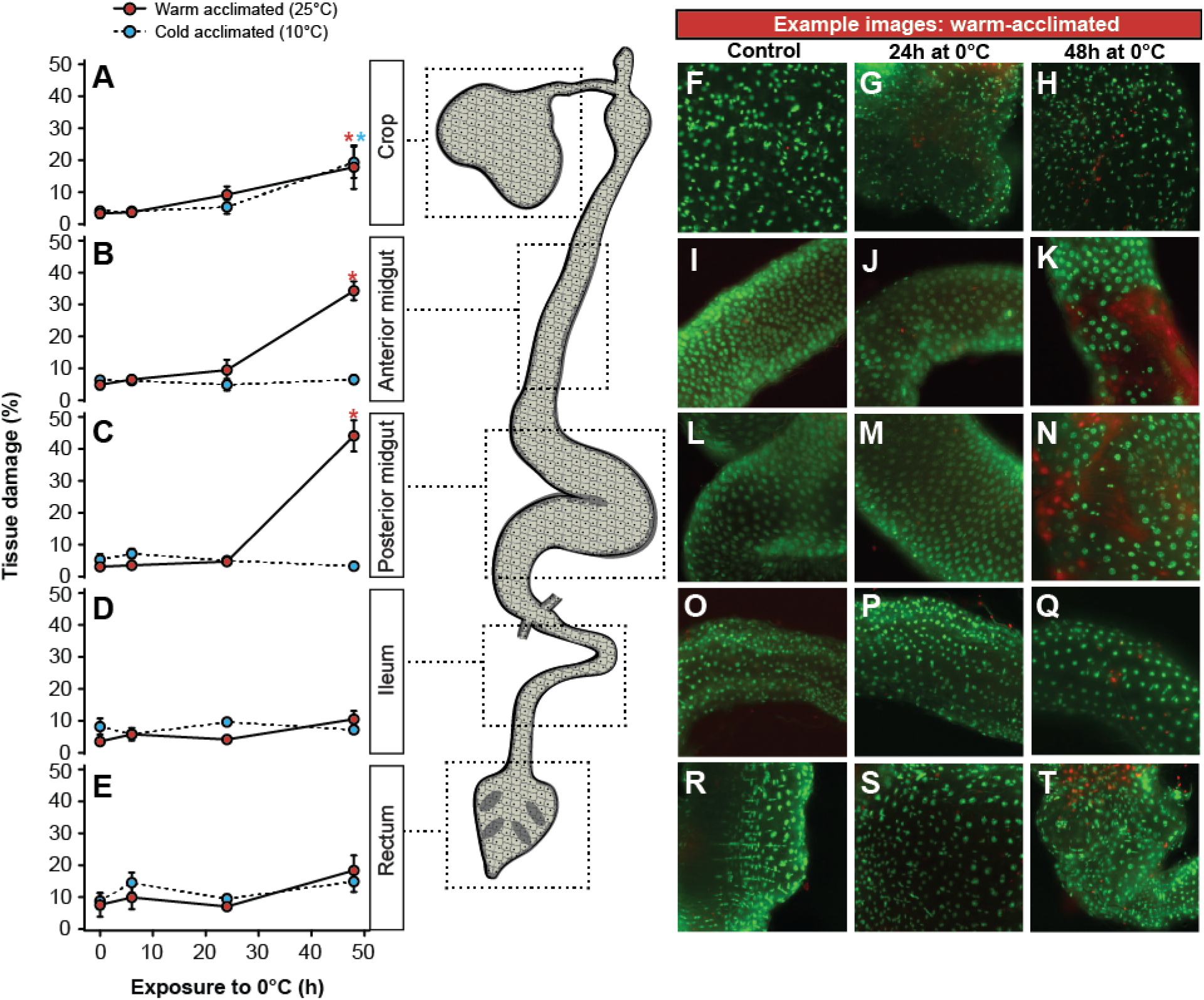
The gut epithelia of warm-acclimated flies suffer tissue damage only after significant paracellular leak has already taken place. The proportion of cells labeled as damaged using the sybr-14 and propridium iodide method for the crop (A), anterior midgut (B), posterior midgut (C), ileum (D) and rectum (E) are shown for both cold-(blue) and warm-acclimated flies (mean ± sem shown). No significant damage was observed in either acclimation group after up to 24h at 0° C (the maximum time at which leak measurements were made), but damage was observed in the crop of both acclimation groups and the anterior and posterior midgut of warm acclimated flies after 48h at 0° C. Asterisks indicate significant increases in tissue damage relative to control (i.e. not cold exposed) samples based on false-discovery rate corrected pairwise t-tests. Error bars that are not visible are obscured by the symbols. Panels F-T show representative examples of tissue staining from warm-acclimated flies after 0 (control), 24, and 48h for the crop (F-H), anterior midgut (I-K), posterior midgut (L-N), ileum (O-Q), and rectum (R-T).

### Junction protein abundance

To determine whether changes in paracellular leak were associated with changes in SJ protein abundance, we tested for differences in the abundance of five SJ proteins in whole-animal homogenates of warm-and cold-acclimated flies (Fig. 4). Coracle (t_5_ = 6.2, *P* = 0.002; Fig. 4A) and Discs-large (t_7_ = 4.0, *P* = 0.005; Fig. 4B) were both significantly more abundant in cold-acclimated flies, by ∼4.5× and 2×, respectively. We observed two distinct bands in our blot for Kune-kune similar in size (at 26 and 72 kDa) to those described in the anal papillae of *Aedes aegypti* ^48^. Neither of these proteins were differentially expressed with thermal acclimation (26 kDa: t_8_ = 0.3, *P* = 0.759; 72 kDa: t_8_ = 0.2, *P* = 0.850; Fig. 4C). Similarly, abundance of Scribble was not significantly affected by cold acclimation (t_6_ = 0.9, *P* = 0.391; Fig. 4D). Mesh, a smooth SJ-associated protein, was significantly lower in cold-acclimated flies relative to warm-acclimated flies (t_7_ = 3.5, *P* = 0.009; Fig. 4E).

**Figure 4.**
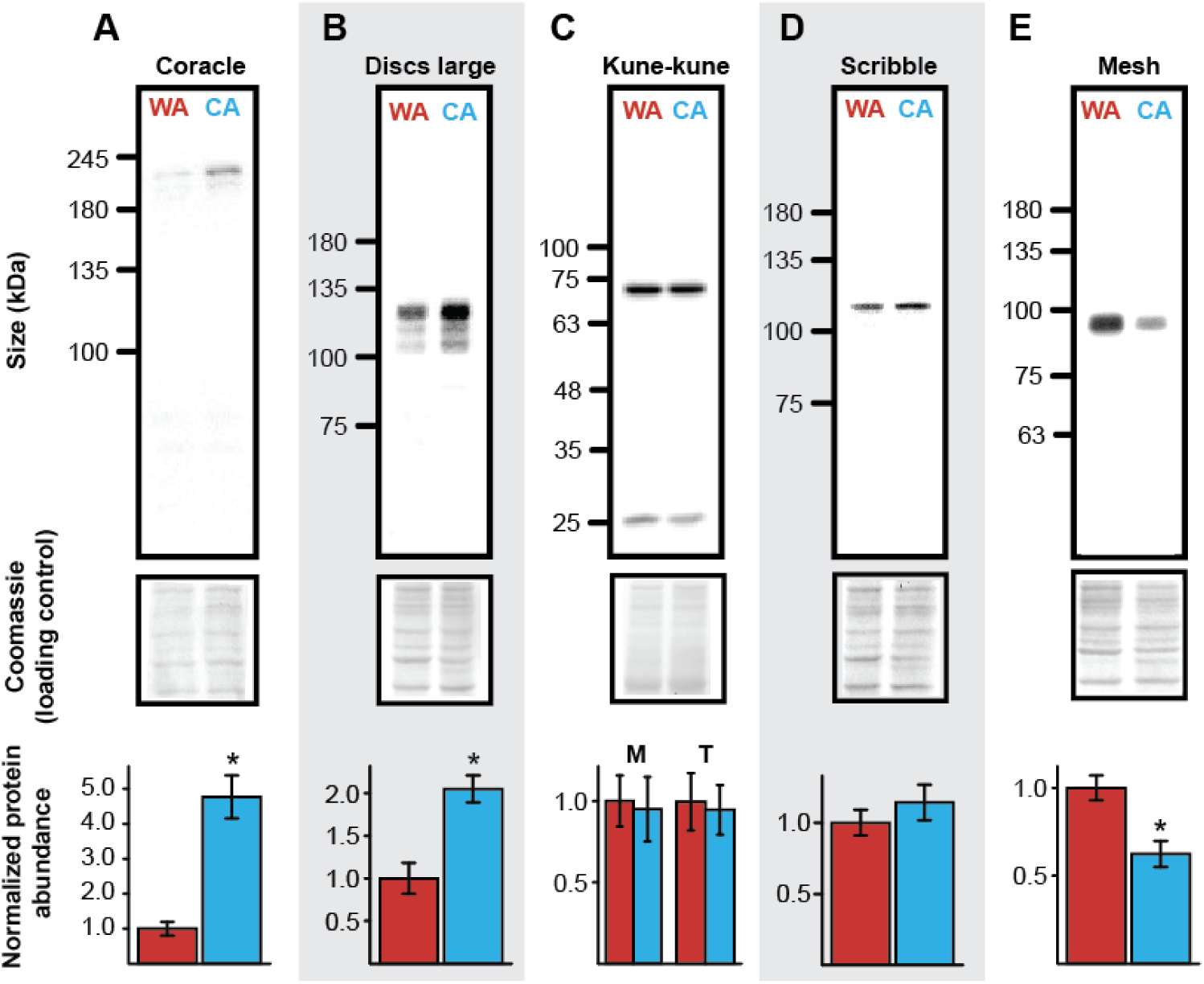
Cold acclimation alters the relative abundance of select septate junction proteins. Protein abundance of A) Coracle, B) Discs large, C) Kune-kune, D) Scribble, and E) Mesh in whole-body samples of female *Drosophila melanogaster* acclimated to warm (WA; 25° C; red) or cold (CA; 10° C; blue) conditions as adults. Protein abundance measured as density (lower panels, mean ± sem) was normalized using a coomassie blue total protein staining method (middle panels). Two bands (at approximately 26 and 72 kDa) were evident for Kune-kune, and these were hypothesized to represent a monomer (M) and trimer (T) form. Asterisks indicate a significant difference in normalized protein expression based on a t-test of log-transformed relative density.

### Midgut junction structure

We used transmission electron microscopy to image regions of cell-cell contact between posterior midgut epithelial cells. The subapical intercellular region was found to be particularly convoluted in cold-acclimated flies (Fig 5A-C), so this was examined quantitatively by measuring the length of the visible intercellular spaces. Cold-acclimated flies had significantly longer regions of cell-cell contact in the subapical intercellular space (F_1,28_ = 14.2, *P* < 0.001; Fig. 5B-D) and this region also contained more visible SJs in cold-acclimated flies (F_1,28_ = 9.4, *P* 0.005; Fig. 5E).

**Figure 5.**
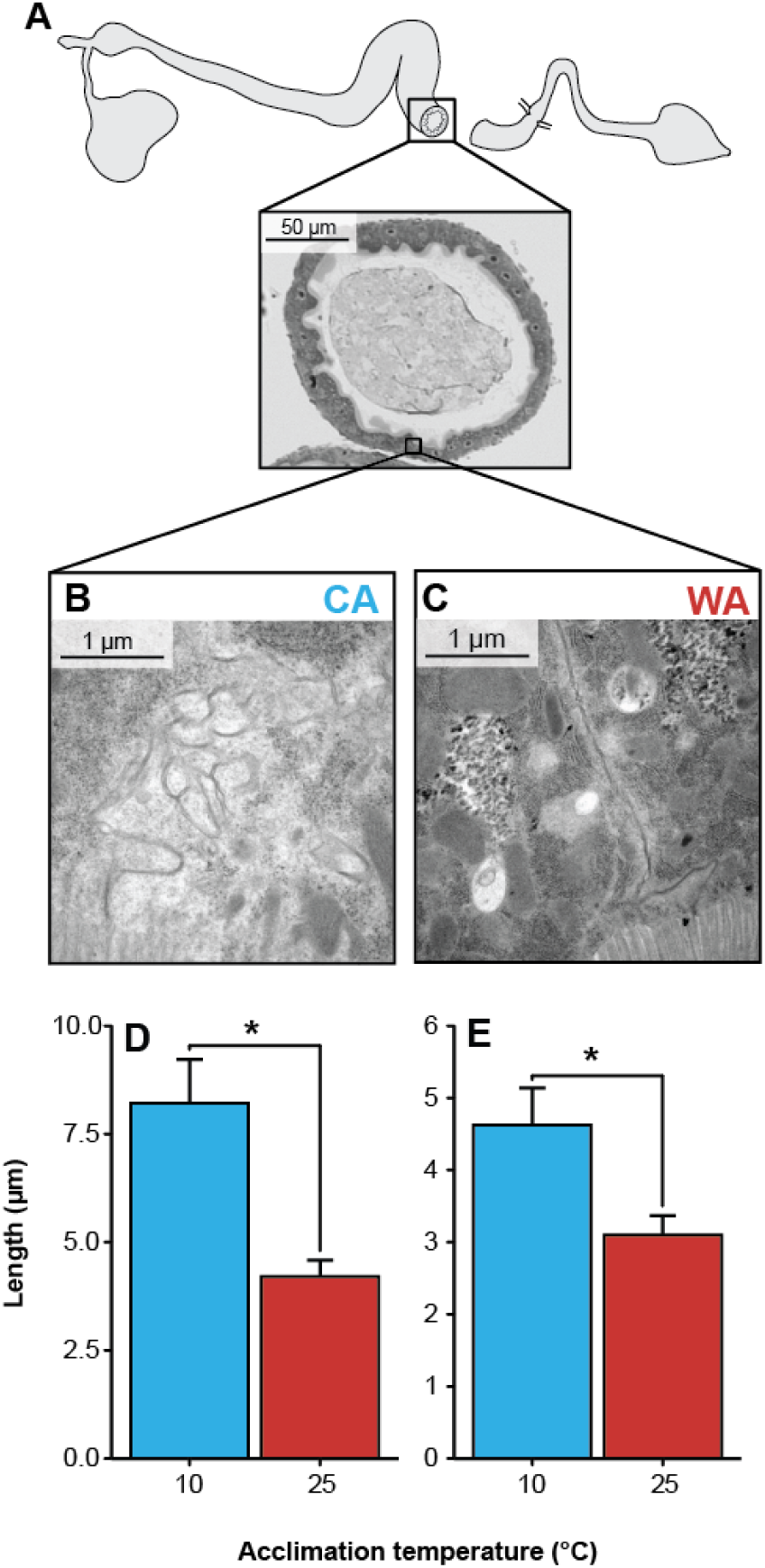
Cold acclimation alters apical cell-cell contact regions in the posterior midgut of *Drosophila melanogaster*. Cross sections from the posterior midgut (alkaline region; A) were visualized by transmission electron microscopy. Representative images of apical regions of cell-cell contact in (B) cold-and (C) warm-acclimated flies. Note the characteristic appearance of the apical intercellular space in warm-acclimated flies (C) in contrast to the longer and more convoluted region of contact between epithelial cells of cold-acclimated flies (B). D) Mean ± sem length of cell-cell contact and E) length of visible septate junctions (SJs) in cold (blue) and warm-acclimated flies (red). Cold acclimated flies had significantly more contact between adjacent cells (*P* <0.0001) and more visible SJs within that region (*P* = 0.005).

## Discussion

### Overview

It is well documented that cold exposure results in a disruption of ion and water balance in insects and that this occurs in conjunction with a decrease in ionomotive enzyme activity^1^. Much of this disruption results from the net leak of Na^+^ and water from the hemolymph into the gut. In this study we demonstrate that cold exposure causes the gut of an insect to leak its contents into the hemolymph through the paracellular pathway. To our knowledge, this is the first evidence of cold-induced epithelial barrier disruption and SJ thermal plasticity in an invertebrate. Epithelial barrier function appears highly relevant to chill tolerance.

### The cold-induced loss of barrier function of gut epithelia

After 24h of feeding on the FITC-dextran diet, cold-acclimated flies exhibited a 7.6-fold lower concentration of labelled dextran in the hemolymph than warm acclimated flies, indicating a reduction in the leak of the paracellular permeability marker across gut epithelia at the acclimation temperature (Fig. 2). During cold exposure, leak of FITC-dextran appeared linear, meaning chilling causes progressive equilibration of this large molecule akin to what is observed for ions leaking across the gut epithelia during chilling^13,49,50^. Notably, progressive leak of dextran also occurred for cold-acclimated flies under the same conditions (Fig. 2C). Cold-acclimated flies are thus not able to completely avoid this problem at 0° C, but instead, alleviate it presumably through changes in their physiology before any cold stress is applied. This result is consistent with the effects of cold-acclimation on ion and water balance in *Drosophila*; acclimation can reduce the rate at which solutes and water leak down their concentration gradients, but cannot completely eliminate these problems at temperatures as low as 0° C. Nevertheless, acclimation can delay injury and death from a cold exposure^20,23^.

### Loss of barrier function occurs while epithelial cells are viable

Chilling can ultimately lead to cell death in a variety of insect tissues^51,52^, which for locust muscle has been linked to depolarization of cell membrane potential as a result of rising hemolymph K^+^ concentrations^15,16^. Accordingly, we sought to examine whether the disruption of barrier function of the gut was driven by cell death or by augmented paracellular leak through an otherwise intact epithelium. Chilling eventually caused cell death, particularly in the midgut, but cell death was coincident with organismal injury and occurred after the times at which loss of barrier function was measured with the FITC dextran assay. Thus, the observed cold-induced loss of barrier function appears to occur prior to epithelial cell damage, and therefore is likely to be driven by changes at the paracellular pathway. This interpretation is supported by the observation that flies had high levels of FITC-dextran in their hemolymph even if they were confirmed to have not suffered any chilling injury (Fig. 2C). Nonetheless, any damage to the gut that does occur later is likely to exacerbate barrier failure.

### Plasticity of barrier function coincides with changes in SJ protein abundance and ultrastructure of apical regions of cell to cell contact

Our results indicate that cold acclimation induced adjustments in paracellular barrier function and these observations would seem to be consistent with a recent study reporting that transcripts encoding ∼75% of known or putative *D. melanogaster* SJ-associated genes were upregulated with cold acclimation (see MacMillan et al., 2016) (Table S1). The correct identification of SJ components that contribute to the observed alterations in paracellular barrier function requires measurement of protein abundance. Hence we began by measuring the relative abundance of five SJ proteins for which we had antibodies readily available in the laboratory. Coracle^53^ is a scaffolding FERM-domain protein and Discs large^54^ is a MAGUK family protein. Both are found in pleated and smooth SJs in *Drosophila*. Scribble is a multi-PDZ and leucine-rich repeat protein found in pleated SJs^55^. Kune-kune is a transmembrane component of pleated SJs and a member of the claudin family of proteins^56^, and Mesh is a protein specific to smooth SJs^57^. The relative abundance of three of these five proteins was altered with cold acclimation (Fig. 4).

Increased abundance of Coracle and Discs large was observed, while there was a decrease in Mesh abundance with cold-acclimation. The results are notable because 1) Coracle, Discs large and Mesh are all found in smooth SJs like those of the midgut, 2) all three proteins are localized to the subapical region of adjacent epithelial cells, and 3) all three proteins are required for paracellular barrier function^53,54,57,58^. The observed changes in protein abundance thus suggest that cold-acclimation induces changes in proteins of apical smooth SJs that are relevant to the barrier function.

Protein and mRNA abundance do not always tightly correlate, largely because of the regulation of post transcription, translation and degradation processes^59^. The increased abundance of Discs large protein mirrors the change in the abundance of *dlg* mRNA observed previously with cold-acclimation (MacMillan et al., 2016; Table S1), but *cora* mRNA abundance did not differ between warm-and cold-acclimated flies (Table S1). This is despite the fact that Coracle protein showed the largest increase in abundance of the five proteins examined. The observation that ∼75% of SJ-associated genes were upregulated with cold acclimation rightly highlighted the SJs as being thermally plastic, but correct identification of specific gene and protein targets clearly requires measurement of realized protein abundance. If Discs large and Coracle (and down regulation of Mesh) are critical to chill tolerance plasticity, knockdown or overexpression of these proteins, particularly in the midgut epithelia, are expected to alter organismal chill tolerance.

Based on these results the ultrastructure of cell-cell contacts between the midgut epithelial cells were examined by electron microscopy and notable differences were observed in the intercellular space of the apical and subapical region. When compared to warm-acclimated flies, the cold-acclimated flies had longer regions of apical/subapical cell-cell contact that appeared convoluted in three dimensions (Fig. 6). These regions also contained more SJs than the same regions in warm-acclimated flies. We hypothesized that this change in apical/subapical cell-cell contact and junctional abundance functions to increase the diffusion distance and epithelial resistance and thereby slow rates of solute and/or solvent leak between the gut lumen and hemolymph in cold-acclimated flies. In other invertebrates as well as vertebrates, alterations in environmental conditions have also been associated with ultrastructural changes in occluding junctions that can be associated with either mitigated or accelerated solute and solvent flux^60–64^. For example, the size and complexity of SJs in the epithelium of snail kidney change in response to alterations in the salinity of the external medium^63^. In low-salinity or hyposmotic medium, where high transepithelial fluid transport is expected, the SJs of snail kidney epithelium have fewer septa and wider intercellular spaces^63^. In contrast, in an isosmotic medium, the SJs of this epithelium become long and have numerous densely packed septa, suggesting reduced transepithelial fluid transport under higher salinity conditions (Khan and Saleuddin, 1981).

## Conclusions

Our results demonstrate that cold exposure perturbs the paracellular barrier properties of the *Drosophila* midgut epithelium and that cold acclimation can partially alleviate this disruption. Cold acclimated flies exhibit profoundly noticeable changes in the ultrastructure of apical and subapical regions of cell-cell contact which coincide with changes in the abundance of select SJ proteins that are known to localize to these regions. Taken together, these results implicate the reorganization or restructuring of SJs in cold acclimation in *Drosophila* midgut. Furthermore, the present study provides a foundation to address the role of paracellular barriers in insect cold tolerance. Perhaps most critical is whether this loss of barrier function specifically facilitates leak of inorganic ions (e.g. Na^+^ and K^+^) or water down their concentration gradients, given that this phenomenon has been repeatedly observed in a wide variety of insects in the cold. A further outstanding question is whether loss of barrier function is specific to the gut or whether other epithelial barriers such as the blood-brain barrier (BBB) are similarly disrupted by chilling and whether thermal acclimation can mitigate such effects. In pigs, for example, mild hypothermia has recently been shown to induce plastic changes in TJ claudin abundance that preserve BBB barrier function during recovery from cardiac arrest^65^. Although here we associated non-freezing chilling injury with barrier disruption, it also remains to be determined whether more cold-hardy insect species suffer similar barrier disruption in the cold or have evolved mechanisms to evade this physiological obstacle.

## Acknowledgements

The authors thank Doug Holmyard (Advanced Bioimaging Centre, Sick Kids^®^ Hospital Research Institute, Toronto, ON, Canada) for assistance with the electron microscopy; Jean-Paul Paluzzi and Magdalena Jaklewicz (York University, Toronto, Canada) for microscope access and support; Prof. Mikio Furuse (National Institute for Physiological Sciences, Okazaki, Aichi, Japan) for generously donating the Mesh antibody.

## Author Contributions

HAM and AD conceived of the study. HAM, GY and SPK collected the data. HAM, GY and SJ analyzed the data. HAM drafted the manuscript, and all authors edited the manuscript.

## Funding

This work was supported by Natural Sciences and Engineering Research Council of Canada (NSERC) Discovery grants to AD and SPK, an NSERC Banting Postdoctoral Fellowship to HAM, and an Ontario Graduate Scholarship to SJ.

